## Supplemental figures for "Heat stress impairs centromere structure and segregation of meiotic chromosomes in Arabidopsis"

This file includes figure supplements

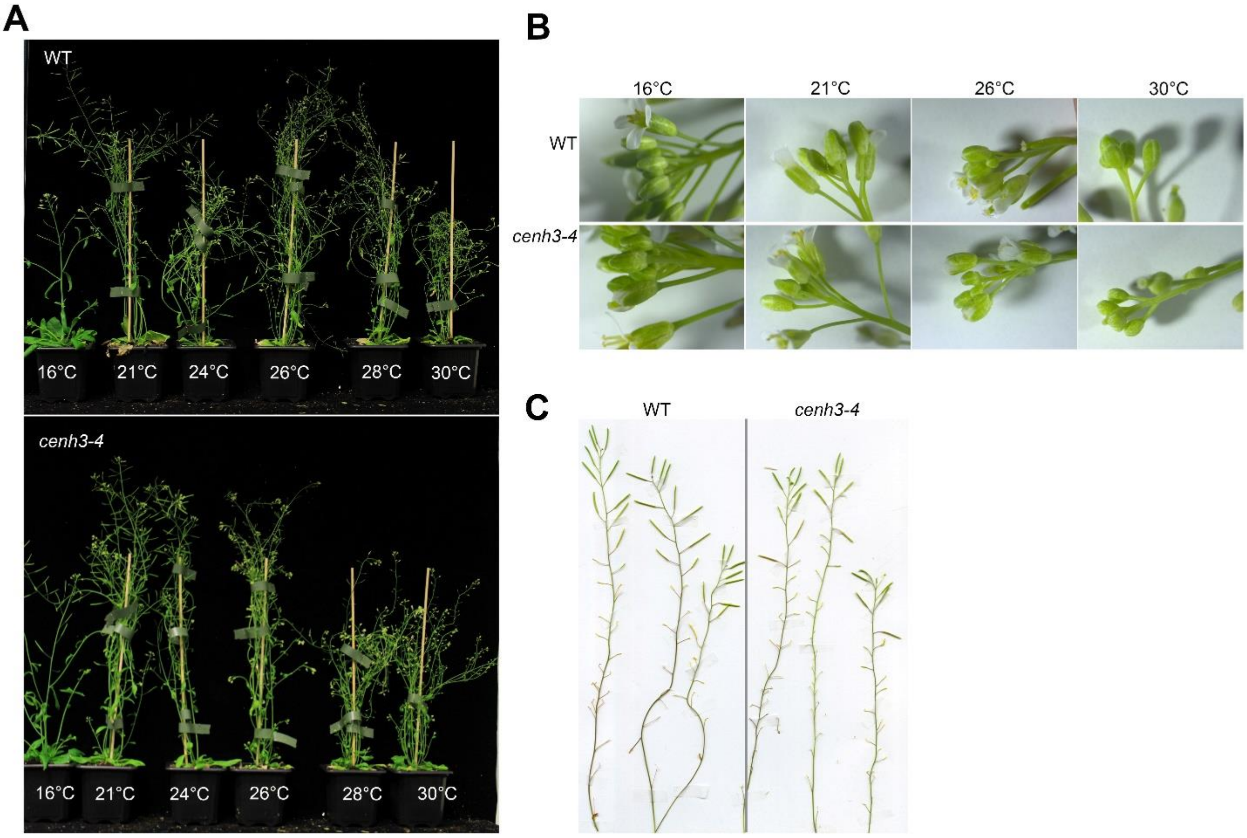


**Figure 1-figure supplement 1. Effect of temperature on growth of wild type and *cenh3-4* plants**.

(A) Approximately six-weeks-old wild type (WT) and *cenh3-4* plants grown at 16, 21, 24, 26, 28 and 30°C. (B) Effect of temperature on WT and *cenh3-4* inflorescence morphology. (C) Restoration of fertility in heat-induced sterile wild type (WT) and cenh3-4 plants; plants were grown at 30°C for 3 weeks and then transferred to 21°C for two more weeks.

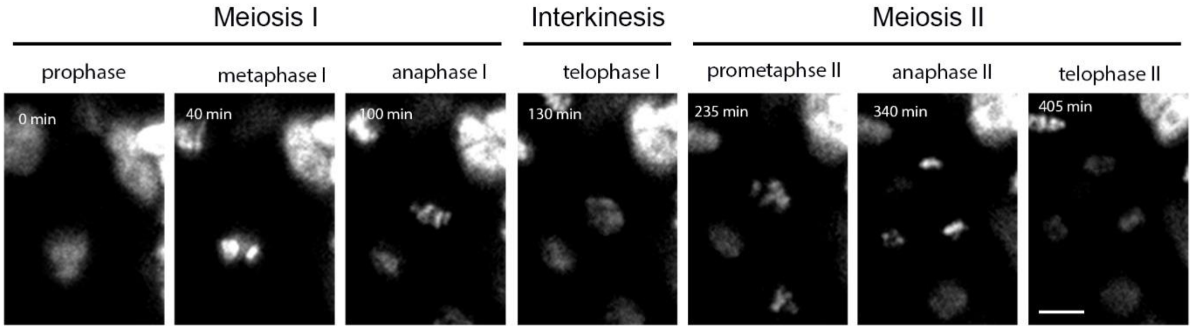


**Figure 2-figure supplement 1.** An example of a time lapse series of HTA10:RFP meiocytes indicating the meiotic stages used for calculating the duration of meiosis. Scale bar=5µm.

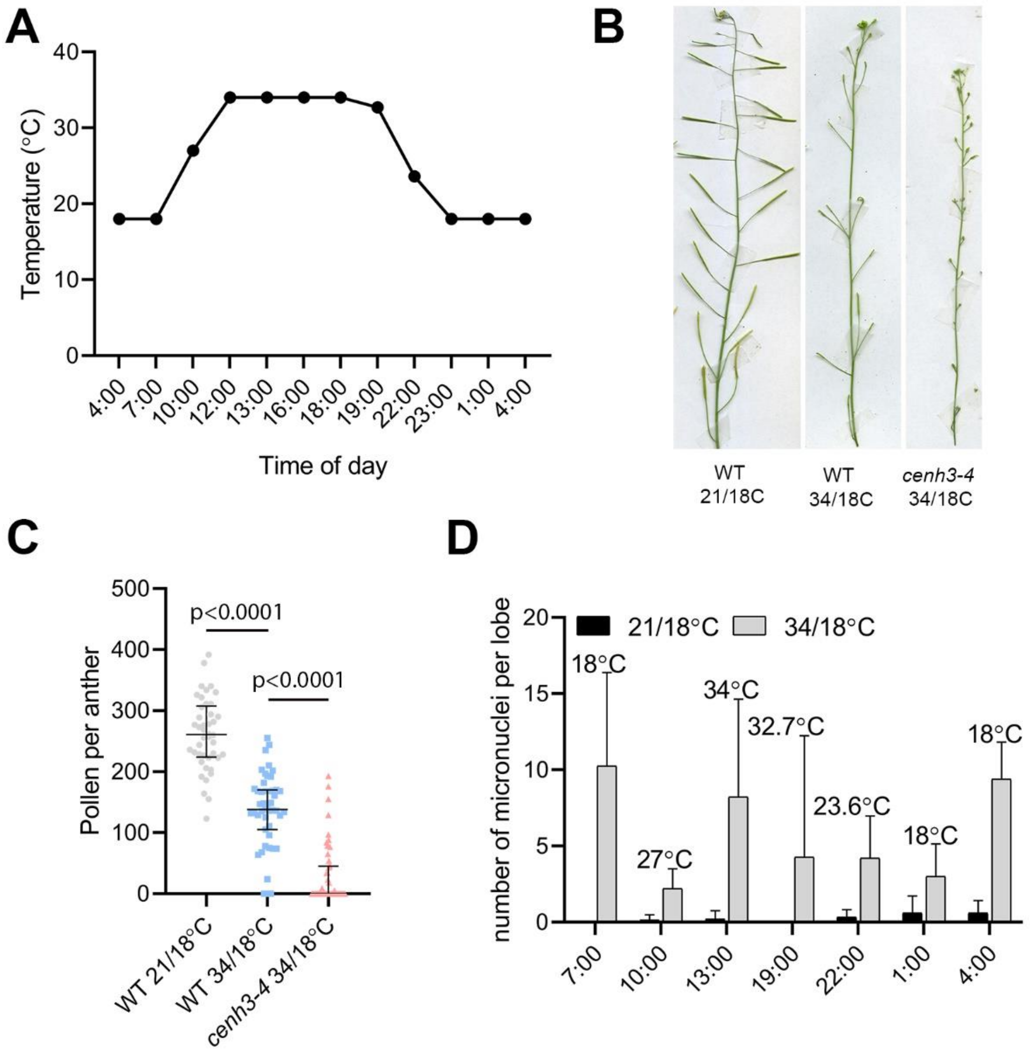


**Figure 2-figure supplement 2. Effect of changing night-day temperatures on wild type and *cenh3-4* plants.**

(A) Temperature distribution in the growth chamber during a 34°C day/ 18°C night regime. (B) Effect of 21°C/ 18°C and 34°C/ 18°C; day/night, temperature regimes on wild type (WT) and *cenh3-4* fertility. (C) Number of viable pollen in wild type at 21°C/ 18°C (n=44), at 34°C/ 18°C (n=47) and *cenh3-4* mutant at 34°C/ 18°C (n=47). (D) Number of produced micronuclei per lobe in wild type anthers of plants grown at 21°C/ 18°C or 34°C/ 18°C. Material was collected at different time points during the day. n=10-13. Error bars depict standard deviation. Source values for (C) and (D) are available in Source data 1.

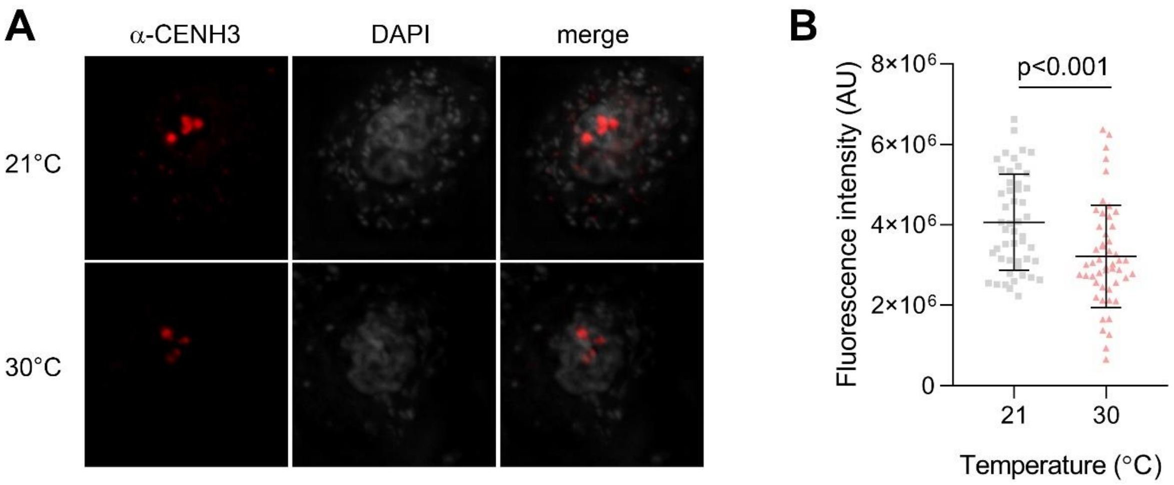


**Figure 3-figure supplement 1. Immunodetection of CENH3 on meiotic chromosomes**

(A) Immunolocalization of CENH3 (red) on pachytene chromosomes stained by DAPI (gray) in PMCs from wild type flowers exposed to 21°C and 30°C. Scale bar=2 µm. (B) Quantification of the CENH3 signal intensity per pachytene centromere in plants grown at 21°C and 30°C. Significance of the difference assessed by the two-tailed t-test is indicated. Source values are available in Source data 1.

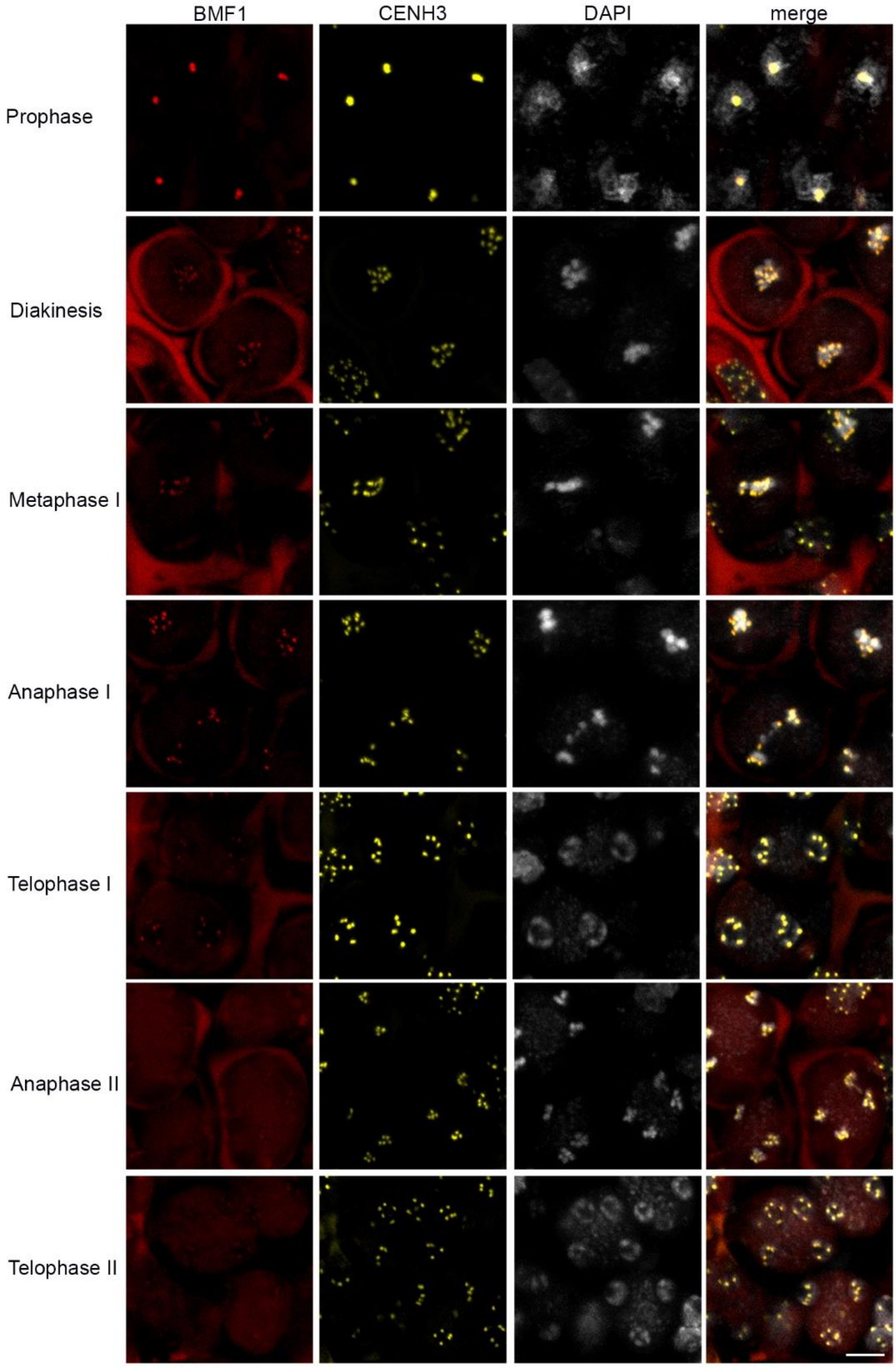


**Figure 3-figure supplement 2.** Colocalization of the kinetochore BMF1:RFP (red) and centromeric eYFP:CENH3 (yellow) proteins through meiosis in DAPI stained meiocytes. Scale bar=5µm.

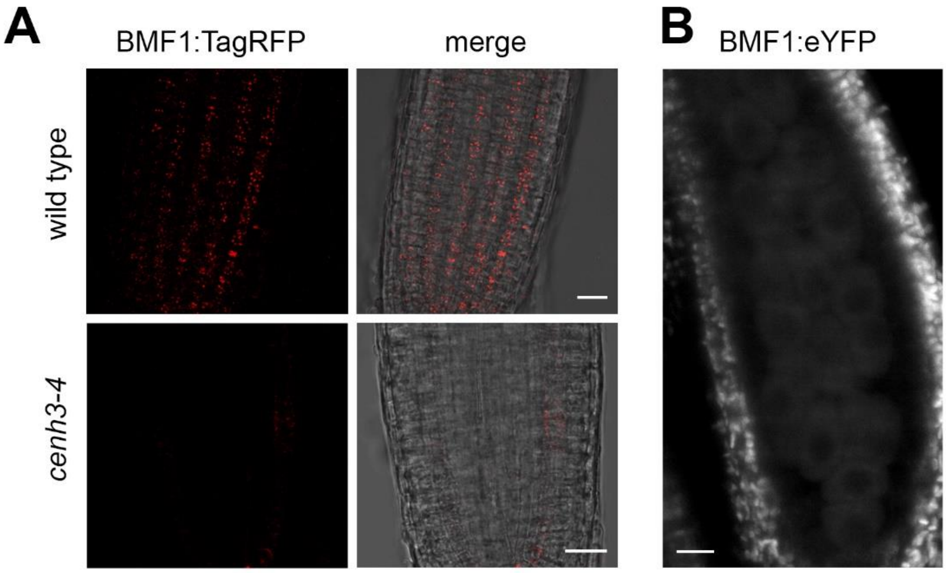


**Figure 3-figure supplement 3. Association of BMF1 with centromeres in *cenh3-4* plants**

(A) BMF1:TagRFP signal in roots of wild type and *cenh3-4* mutants. The same conditions of image acquisitions were used. Scale bar = 20 µm. (B) No BMF1:eYFP signal is detectable in PMCs of *cenh3-4* mutants grown at 21°C even with the maximum intensity of the excitation laser. Scale bar = 20 µm.

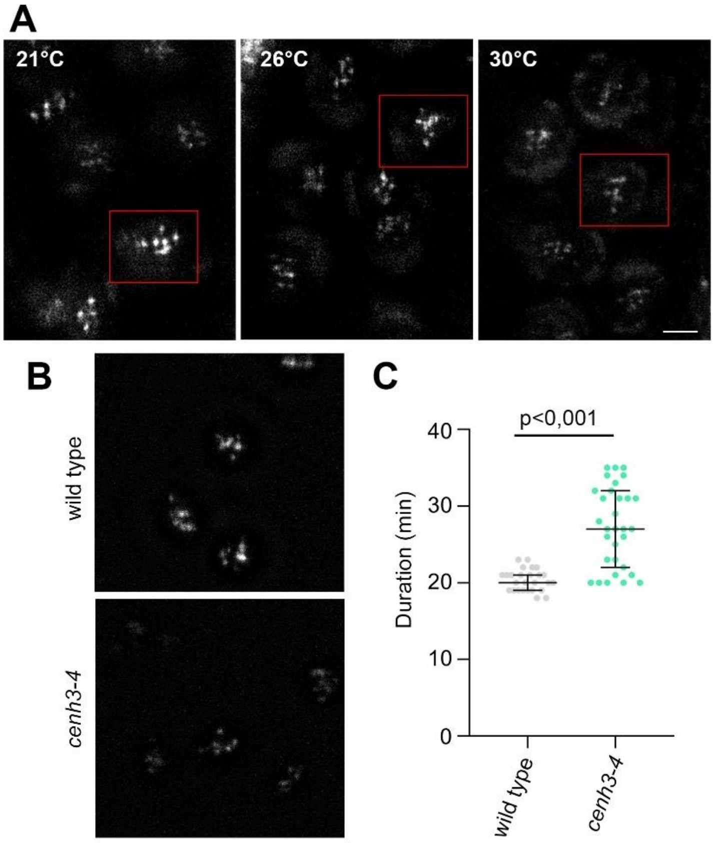


**Figure 4-figure supplement 1. Association of BMF3 with centromeres in *cenh3-4* plants**

(A) BMF3:GFP signal in meiosis I obtained by live cell imaging in PMCs incubated at indicated temperature. Intensity of the BMF3:GFP signal changes in the course of meiosis and the red rectangles indicate PMCs with the peak BMF3:GFP signal intensity. Scale bar = 5 µm. (B) BMF3:GFP signal in meiosis I in wild type and *cenh3-4* plants obtained by live cell imaging microscopy. Scale bar = 5 µm. (C) Duration of the BMF3:GFP signal in PMCs during meiosis I in wild type and *cenh3-4* plants. Each datapoint corresponds to one meiocyte. Significance of the difference assessed by two-tailed t-test is indicated. Source values are available in Source data 1.
